# Cryptic diversity in the model fern genus *Ceratopteris* (Pteridaceae)

**DOI:** 10.1101/2020.02.12.946202

**Authors:** Sylvia P. Kinosian, William D. Pearse, Paul G. Wolf

## Abstract

Cryptic species are present throughout the tree of life. They are especially prevalent in ferns, because of processes such hybridization, polyploidy, and reticulate evolution. In addition, the morphological simple body plan of ferns limits phenotypic variation and makes it difficult to detect crypic species in ferns without molecular work. The model fern genus *Ceratopteris* has long been suspected to harbor cryptic diversity, specifically in the highly polymorphic *C. thalictroides*. Yet no studies have included samples from throughout the pan-tropical range of *Ceratopteris* or utilized genomic sequencing, making it difficult to assess the full extent of cryptic variation within this genus. Here, we present the first multilocus phylogeny of the genus using reduced representation genomic sequencing (RADseq) and examine population structure, phylogenetic relationships, and ploidy level variation. We recover similar species relationships found in previous studies, find support for a named cryptic species as genetically distinct, and identify a novel putative species from within *C. thalictroides sensu latu* in Central and South America.

## 1. Introduction

Describing the formation of species is essential to understanding the patterns and processes that create biodiversity. Detecting differences among species becomes increasingly challenging when taxa are morphologically similar, but genetically distinct and reproductively isolated (Bickford et al. 2007; Masuyama 1992; Paris et al. 1989). Such cryptic species have often been historically described as one larger species, or species complex, due to morphological similarities (Paris et al. 1989). However, advances in molecular methods have revealed that cryptic species can be monophyletic with disparate ecological niches and functions (Amato et al. 2007; Hebert et al. 2004; Sattler et al. 2007; Southgate et al. 2019); in contrast, cryptic species complexes can be paraphyletic and separated by considerable evolutionary time, yet look similar and occupy comparable niches (Amor et al. 2014; Cunnington et al. 2005).

Despite the recent shift in molecular approaches to investigate cryptic species, much is still unknown about their evolutionary history and ecosystem functions. Cryptic species complexes often have very broad distributions (Der et al. 2009; Knowlton 1993; Nygren 2014) and can occur in sympatry, allopatry, or parapatry (Bickford et al. 2007), making adequate sampling challenging. In ferns, deciphering species boundaries can be even more difficult due to their ease of dispersal via spores, making gene flow possible across vast distances (Barrington 1993; Tryon 1970). Compared to seed plants, ferns have a very high prevalence of polyploidy and reticulate evolution (Barrington et al. 1989; Paris et al. 1989; Otto and Whitton 2000; Otto and Whitton 2000). Such genomic changes can contribute to cryptic variation by altering niche requirements or offspring viability (Otto 2007; Southgate et al. 2019; Masuyama et al. 2002), yet be essentially phenotypically invisible (Patel et al. 2019).

The fern genus *Ceratopteris* Brong. is a pan-tropical aquatic clade consisting of seven named species, three of which are cryptic and tetraploid (PPGI 2016; Lloyd 1974; Masuyama and Watano 2010). *Ceratopteris* is perhaps best known for the model organism *C. richardii*, which has been used as such since the late twentieth century (Banks 1994; Hickok et al. 1987; Hickok et al. 1995). Sometimes called the *“Arabidopsis* of the fern world” (Sessa et al. 2014), *C. richardii* is an ideal model system because of its fast life cycle and ease of cultivation (Hickok et al. 1987); in addition, it can be transformed with recombinant DNA (Muthukumar et al. 2013; Plackett et al. 2014), has a reference ontogeny framework (Conway and Di Stilio 2019), and is currently the only homosporous fern to have a whole genome sequence (Marchant et al. 2019). In addition to *C. richardii*, other *Ceratopteris* species have been studied in a lab environment (Hickok and Klekowski 1974; Hickok 1977), and have potential for further research. Concerningly, the species boundaries within the genus are blurry, and there is evidence that all species in the group (both diploid and tetraploid) hybridize to some extent (Adjie et al. 2007; Hickok and Klekowski 1974; Hickok 1977; Hickok 1979; Lloyd 1974). In such a well-utilized model genus, there is still a need to better understand species boundaries, evolutionary history, and occurrences of cryptic species. In particular, we need to understand the evolutionary dynamics of the *Ceratopteris* genus as a whole in order to best utilize this model system for future work.

The first and only comprehensive monograph for *Ceratopteris* was written by Lloyd in 1974, and employed a matrix of morphological traits to identify four species in the genus. However, *Ceratopteris* is a notorious group in terms of morphology: there are relatively few informative physical characters (Lloyd 1974); in addition, habitat and developmental stage can have a large effect on plant phenotype (Masuyama 1992). Most challenging for species delimitation, however, is that rampant hybridization in the genus is known to further alter morphology and so make field identification difficult (Hickok and Klekowski 1974; Lloyd 1974; Masuyama and Watano 2010). While Lloyd named four species, he noted that *C. thalictroides*, the most widespread species in the genus, is “highly polymorphic” (Lloyd 1974) and is likely a cryptic species complex. In response, Masuyama and colleagues conducted a series of studies on Asian plants under the name of *C. thalictroides*, examining chloroplast DNA and cross-breeding (Masuyama et al. 2002), cytological characteristics (Masuyama and Watano 2005), morphological traits (Masuyama 1992; Masuyama 2008), and nuclear DNA (Adjie et al. 2007). As a result, three cryptic species and two varieties were named from entities originally described as *C. thalictroides* (Masuyama and Watano 2010). Molecular evidence suggests that all three cryptic species of *C. thalictroides* have independent hybrid allopolyploid origins and are paraphyletic (Adjie et al. 2007), the latter providing the most compelling evidence that they cannot be described under the same name. However, this study included only one nuclear marker; multilocus analyses are generally needed to substantiate paraphyly and polyploid origins (Eaton and Ree 2013; Jorgensen and Barrington 2017), and particularly in a case as complex as *C. thalictroides*. In addition, while cryptic species in *Ceratopteris* have been investigated in the Old World, there is a similar problem of morphological variability in New World *C. thalictroides* (Masuyama and Watano 2010; Lloyd 1974), indicating a potential for more cryptic species to be discovered.

As a well-studied, pan-tropical group known to include cryptic species, *Ceratopteris*, and in particular *C. thalictroides*, is an ideal system in which to study the process of cryptic speciation. In this study, we produce the first multilocus genomic analysis of *Ceratopteris* using restriction-site associated DNA sequencing (RADSeq). RADSeq is a cost-effective way to generate a large amount of genomic data, and many downstream analysis tools are currently available. We apply three approaches to identify cryptic species. First, we estimate population structure and hybridization; second, we reconstruct phylogenetic relationships among samples; finally, we investigate ploidy levels across individuals. This study is a step towards resolving species boundaries within this complex genus. Our work will also provide a phylogenetic reference for future studies on *Ceratopteris*, as well as find areas within the genus in need of additional research.

## 2. Materials and methods

To best analyze the cryptic diversity within *Ceratopteris* we gathered samples from across its pan-tropical distribution for 5 of 7 species. We utilized RADSeq to generate a very large genomic dataset, which was processed using the ipyrad pipeline (Eaton and Overcast 2020) for downstream analysis. Our three-part approach for assessing cryptic diversity takes advantage of the flexibility of RADSeq data, and provides a unique window into the processes of speciation and diversification in a complex genus.

All parameter values and code for data processing and downstream analyses can be found on GitHub (github.com/sylviakinosian/ceratopteris_RADSeq).

### 2.1. Taxon sampling

Because of its broad distribution, we chose to use a combination of herbarium and silica-dried material from field collections to sample *Ceratopteris* (Fig. 1). We started with 90 samples, 56 from herbarium specimens and 34 silica-dried tissue. These samples represented five of seven named species of *Ceratopteris*; we did not include *C. oblongiloba* Masuyama & Watano and *C. froesii* Brade. These two missing species were recently named cryptic species of *C. thalictroides* (L.) Brongn. (Masuyama and Watano 2010). We also included one sample of *Acrostichum aureum* from Australia for use as an outgroup. Herbarium specimens were chosen based on age (less than about 30 years old) and color (leaves still green). These specimens were collected from the Harvard University Herbaria (HUH), the Steere Herbarium at the New York Botanic Garden (NY), the University of California, Berkeley (UC), the United States National Herbarium at the Smithsonian (US), and the Pringle Herbarum at the University of Vermont (VT). Fresh tissue collections were obtained from Taiwan, China, Costa Rica, and Australia. All field collections were stored on silica gel and vouchers were deposited at the Intermountain Herbarium (UTC) and James Cook University (JCT). See Supplementary data 1 for full details of specimen sources.

**Figure 1:**
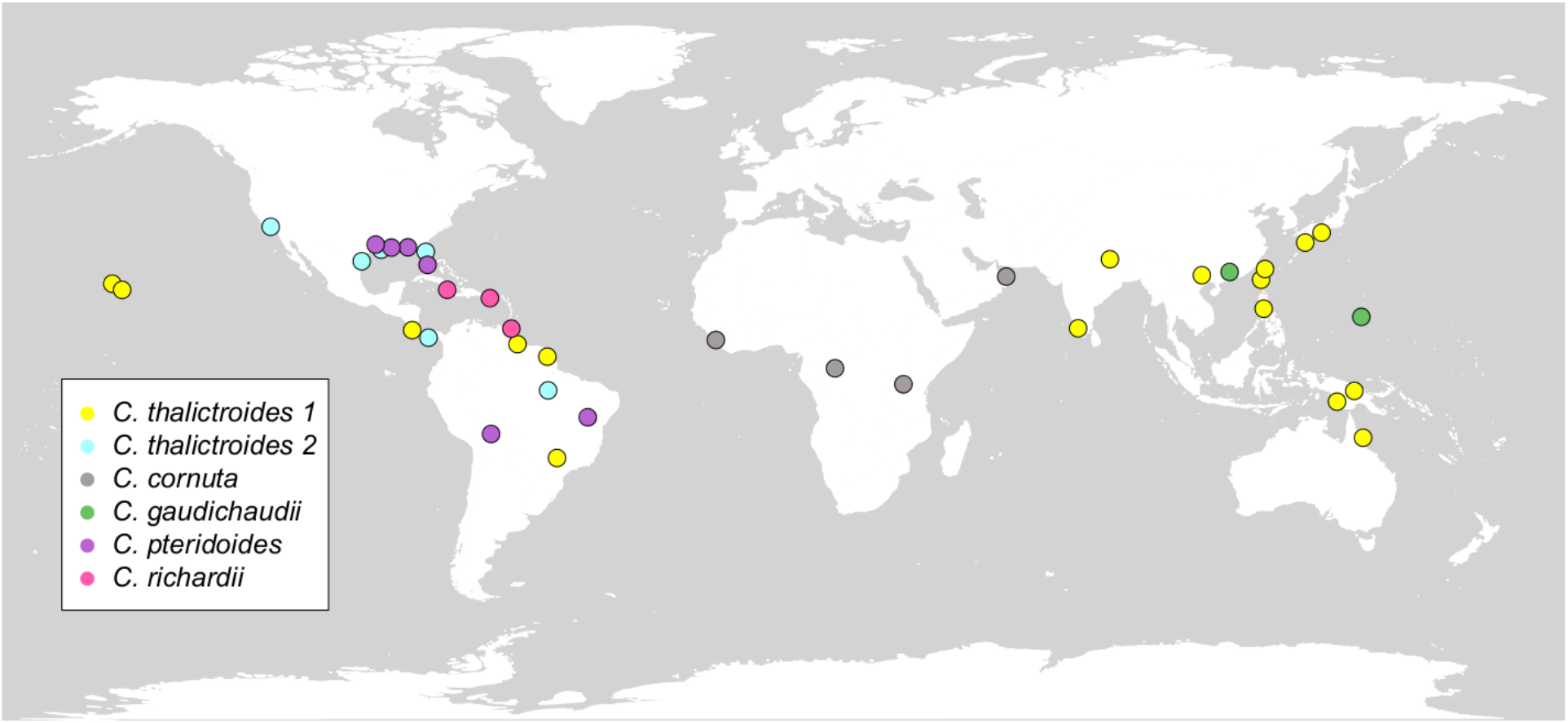
Collection localities for samples of *Ceratopteris* included in the present study. Not all accessions are included on this map because many herbarium specimens did not have GPS coordinates, or several samples were from nearly identical locations.

### 2.2. DNA extraction

Silica-dried plant tissue samples were supplied to the University of Wisconsin-Madison Biotechnology Center. DNA was extracted using the QIAGEN DNeasy mericon 96 QIAcube HT Kit, and quantified using the Quant-iT^TM^ PicoGreen^®^ dsDNA kit (Life Technologies, Grand Island, NY). 17 specimens were extracted using a modified CTAB method (Doyle and Doyle 1987) by SPK at Utah State University. These specimens were send to the University of Wisconsin-Madison Biotechnology Center, analyzed for quality and then pooled with the rest of the samples.

### 2.3. Library construction and sequencing

Libraries were prepared following Elshire *et al*. (2011) with minimal modification; in short, 100 ng of DNA was digested using PstI and BfaI (New England Biolabs, Ipswich, MA) after which barcoded adapters amenable to Illumina sequencing were added by ligation with T4 ligase (New England Biolabs, Ipswich, MA). 96 adapter-ligated samples were pooled and amplified to provide library quantities amenable for sequencing, and adapter dimers were removed by SPRI bead purification. Quality and quantity of the finished libraries were assessed using the Agilent Bioanalyzer High Sensitivity Chip (Agilent Technologies, Inc., Santa Clara, CA) and Qubit^®^ dsDNA HS Assay Kit (Life Technologies, Grand Island, NY), respectively. A size selection was performed to obtain 300 - 450 BP fragments. Sequencing was done on Illumina NovaSeq 6000 2×150 S2. Images were analyzed using the standard Illumina Pipeline, version 1.8.2.

### 2.4. Data processing

Raw data were demultiplexed using stacks v. 2.4 (Catchen et al. 2011; Catchen et al. 2013) process_radtags allowing for a maximum of one mismatch per barcode. Demultiplexed FASTQ files were paired merged using ipyrad version 0.7.30 (Eaton and Overcast 2020). Low quality bases, plus adapters and primers were removed from each read. Filtered reads were clustered at the default setting of 85%, and we required a sequencing depth of 6 or greater per base and a minimum of 30 samples per locus to be included in the final assembly. The ipyrad pipeline defines a locus as a short sequence present across samples. From each locus, ipyrad identifies single nucleotide polymorphisms (SNPs); these SNPs are the variation used in our downstream analyses. Since stacks and ipyrad assume diploidy, we processed all samples as such. However, some species of *Ceratopteris* are non-diploid (Adjie et al. 2007; Masuyama and Watano 2010), and so we explore the ploidy of each sample more thoroughly later in our analyses.

Our initial sampling contained 95 specimens of *Ceratopteris* and 1 sample of *Acrostichum aureum*. Of the starting samples of *Ceratopteris*, 26 either extracted very poorly or yielded few loci (less than 1000 loci per individual). Those samples were removed from downstream analysis. We also removed 13 samples from Australia, an area we were able to sample on a very fine spatial scale: one individual per Australian population was included in the final dataset so we did not over represent Australian specimens in our analyses. Our final data set included 56 samples of *Ceratopteris* and 1 sample of *A. aureum*. We included five specimens that were lower quality (see Supplementary data 1) because they were important in achieving a more even sampling distribution. They did not cluster with each other in our analyses, and their assignments in the population structure and phylogenetic analyses were biologically plausible. We did two separate runs of the ipyrad pipeline: one with *A. aureum* for phylogenetic analysis, and one without for ploidy estimation and population structure analysis. Because *A. aureum* is sister to *Ceratopteris* and separated by over 30 million years (PPGI 2016), the total number of SNPs retrieved decreases when *A. aureum* is included.

### 2.5. Population and genomic structure analysis

Because species of *Ceratopteris* are known to hybridize (Hickok and Klekowski 1974), but also form reproductively isolated cryptic species (Masuyama et al. 2002), we wanted to examine population structure of the genus across its distribution. The program STRUCTURE v. 2.3.4 (Pritchard et al. 2000) is designed to determine admixture (hybridization) between populations of one or several closely related taxa. The program assumes that each individual’s genome is a mosaic from *K* source populations and use genotype assignments to infer population structure and admixture. In our analysis, we ran STRUCTURE for *K* = 2-7 with 50 chains for each K. We then used CLUMMPAK (Kopelman et al. 2015) to process the STRUCTURE output and estimate the best *K* values (Evanno et al. 2005; Pritchard et al. 2000).

### 2.6. Species tree inference

To construct a phylogeny for the samples of *Ceratopteris*, we used the program TETRAD, which is included in the ipyrad analysis toolkit and is based on the software SVDQuartets (Chifman and Kubatko 2015). TETRAD utilizes the theory of phylogenetic invariants to infer quartet trees from a SNP alignment. Species relationships are estimated for all quartet combinations of individuals, then all quartet trees are joined into a species tree with the software wQMC (Avni et al. 2015). Quartet methods were designed to reduce computational time for tree-building with maximum likelihood methods (Ranwez and Gascuel 2001), making them particularly useful for SNP datasets such as these.

We ran TETRAD via Python 2.7.16 (www.python.org) for 56 samples of *Ceratopteris*, and 1 sample of *Acrostichum aureum* as the outgroup. Our input consisted of 57 individuals and 29849 SNPs; we ran 100 bootstrap iterations for the final consensus tree. We then plotted the final tree in R using the package phytools (Revell 2012) and custom plotting functions.

### 2.7. Polyploidy analysis

Due to the presence of numerous cryptic species and morphological variation across the genus, we hypothesize there may be cryptic cytotype variation within species, especially the polymorphic *C. thalictroides*. Since a majority our specimens were from herbarium specimens, we could not perform flow cytometry or chromosome squashes. We utilized the R v. 3.5.2 (R Core Team 2018) package gbs2ploidy to detect ploidy variation across samples. This package was designed to detect ploidy levels in variable cytoptype populations (specifically quaking aspen, *Populus tremuloides*) using low-coverage (2X) genotyping-by-sequencing (GBS) or RADSeq data; it was also tested on simulated data, and can detect diploids, triploids, or tetraploids (Gompert and Mock 2017). This package estimates ploidy from allele ratios, which are calculated from genome-average heterozygosity and bi-allelic SNPs isolated from a variant calling format (VCF) file, which is included in the output from ipyrad.

To perform our analyses with gbs2ploidy, we first used the script vcf2hetAlleleDepth.py (github.com/carol-rowe666/vcf2hetAlleleDepth) to convert the VCF file produced by ipyrad to the format needed for gbs2ploidy. We then ran gbs2ploidy, using the function estprops to estimate ploidy for each individual. We plotted the output from gbs2ploidy in R, with the mean posterior probability for allelic ratios on the *y* axis and the 1:1, 2:1, and 3:1 ratios on the *x* axis (see Fig. 2); we also included error bars for the 95% equal tail probability intervals (ETPIs). We assigned ploidy to each individual using the highest posterior mean estimate for a certain allelic proportion. If the ETPIs overlapped, we considered it an ambiguous assignment. We required sequencing depth to be 6X or greater during our data processing, which suggests that the ploidy assignments are likely accurate.

**Figure 2:**
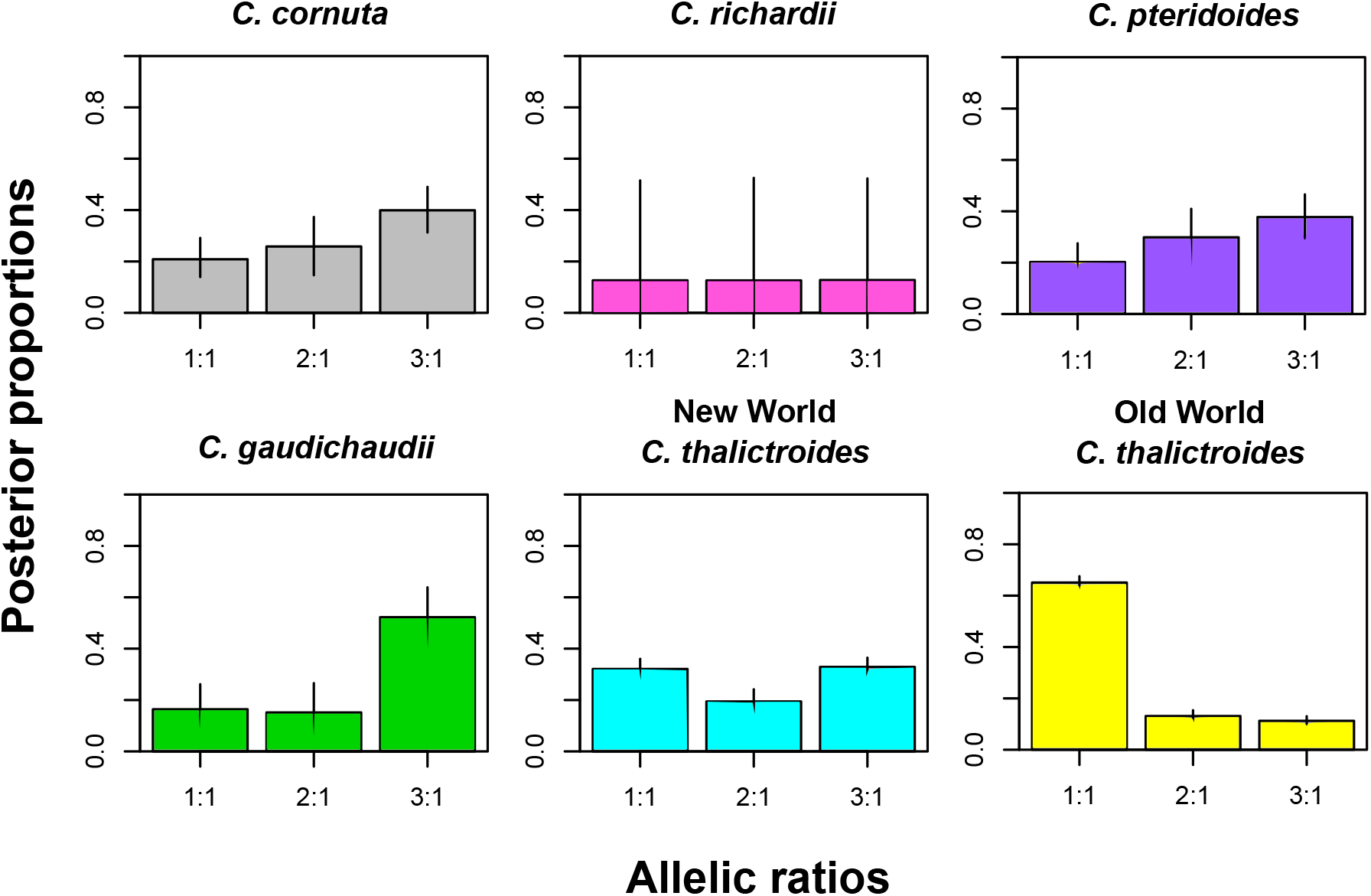
Ploidy estimates for a subset of individuals. The X axis displays the estimated allele ratios (1:1, 2:1, 3:1), and the Y axis shows the Bayesian posterior proportion of allelic ratios; 95% credible intervals are shown on each bar. The allelic ratio with the highest posterior estimate was used to assign ploidy to an individual. If the credible intervals overlapped, ploidy was assigned as ambiguous. Individuals included here are: *Ceratopteris cornuta* (Cr03), *C. gaudichaudii* (Ga02), *C. pteridoides* (Pt03), New World *C. thalictroides* (Th17), and Old World *C. thalictroides* (Th51). See Supplementary data 1 for sample IDs and all ploidy estimates.

## 3. Results

We retrieved an average of 2.58 x 10^6^ raw reads per sample (See Supplementary data 1). From the 60 herbarium samples and 36 silica-dried specimens, we retrieved an average of 5954 and 15029 loci, respectively. On average, we recovered nearly three times the number of loci from silica-dried specimens compared to the herbarium specimens. The 17 herbarium samples had been previously extracted before sending to the University of Wisconsin; the remaining samples (of both herbarium and silica-dried tissue) were extracted from leaf tissue at UW facilities. The UW facilities obtained much higher-quality extracted DNA than we did in our own lab, which contributed to some of the disparity between herbarium and silica-dried DNA quality.

### 3.1 Population and genomic structure analysis

We used the best *K* method by Evanno et al. (2005), which determined a *K* value of 4. The best *K* estimate described in the STRUCTURE manual (Pritchard et al. 2000) determined a *K* value of 5. We also visually examined *K* = 2 - 7 and determined that *K* = 5 was the most biologically meaningful because it separated each named species, and also showed some intra-specific variability (Fig. 3). Increasing *K* past 5 did not add any meaningful population clusters.

**Figure 3:**
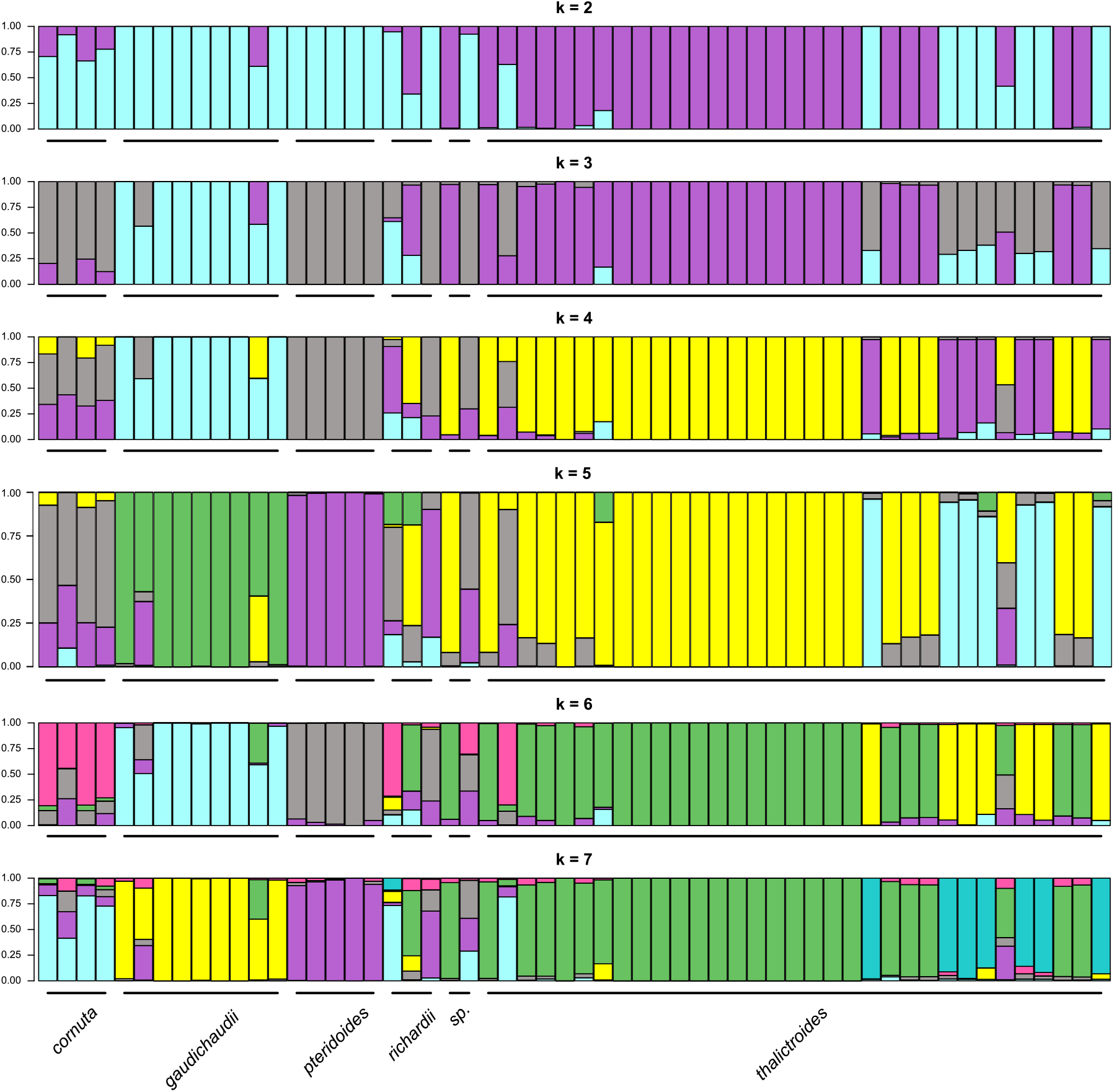
STRUCTURE plots for *K* = 2 - 7. *K* = 5 was determined to be the most biologically informative, as increasing *K* past 5 did not add any biologically meaningful groups. At *K* = 5, all named species of *Ceratopteris* fall into distinct populations, and C. thalictroides is split into three groups: Old World (yellow), New World (blue), and individuals grouping with *C. gaudichaudii* (green).

Named species of *Ceratopteris* mostly clustered together at *K* = 5 (Fig. 3). *Ceratopteris cornuta* and *C. pteridoides* appeared as distinct populations. Unfortunately, all samples of *C. richardii* had very poor sequencing depth and quality, so we cannot be sure if their variable population assignments are due to mis-identification of herbarium specimens, poor data quality, or unknown variation within the species.

*Ceratopteris thalictroides sensu latu* split in to three groups at *K* = 5. The first group consisted of several individuals identified as *C. thalictroides* (on herbarium sheets or in the field) that clustered with *C. gaudichaudii*, a cryptic species of *C. thalictroides* (Masuyama and Watano 2010). The remaining individuals of *C. thalictroides* grouped into two populations, one consisting of mostly Old World individuals, and the other as entirely New World individuals. These two populations were strongly differentiated by STRUCTURE, being separated at all *K* values.

### 3.2. Species tree inference

The phylogeny resulting from our species quartet inference shows a similar pattern to the STRUCTURE output (Fig. 4). Most individuals of *Ceratopteris pteridoides* form a monophyletic group, with one individual sister to the remaining species. *Ceratopteris cornuta* came out in two places in the phylogeny: sister to Old World *C. thalictroides*, and in a grade outside the *C. gaudichaudii*, Old World and New World *C. thalictroides* clade. Our phylogeny grouped all individuals placed in the *C. gaudichaudii* population by STRUCTURE in a monophyletic clade, sister to New World *C. thalictroides*. Samples of *C. richardii* were also found in multiple places in the phylogeny; however, due to their lower data quality this is not surprising and little inference can be made about its phylogenetic position.

**Figure 4:**
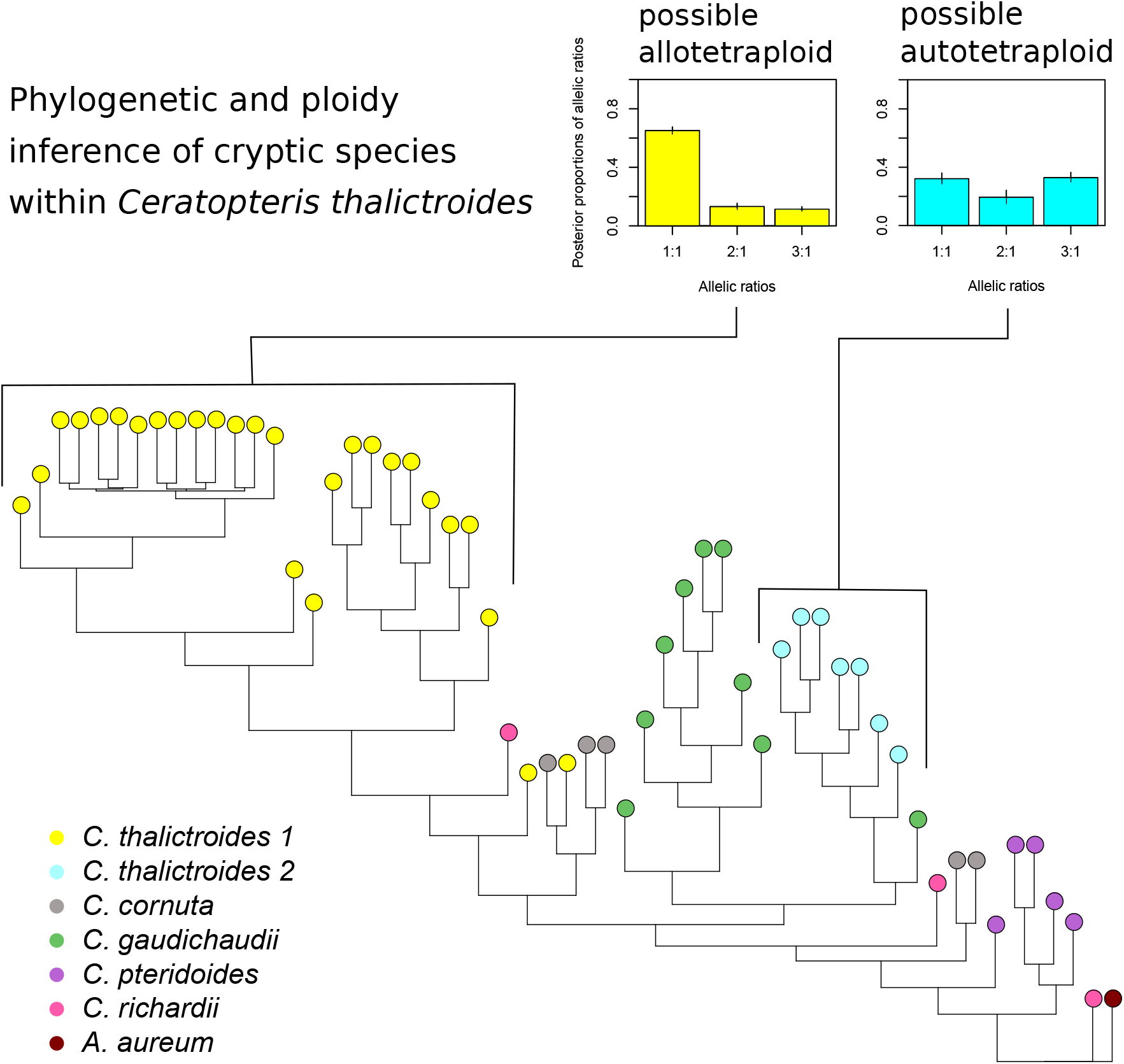
Phylogeny of *Ceratopteris* generated via quartet methods, and ploidy assignments for the Old and New World *Ceratopteris thalictroides* clades.

### 3.3. Ploidy assignment across species

*Ceratopteris cornuta, C. pteridoides*, and *C. richardii* are known as a diploid (n = 39) (Adjie et al. 2007; Hickok 1977); *Ceratopteris thalictroides* and *C. gaudichaudii* are known as tetraploid, with some cytotypic variation (n = 77, 78) (Adjie et al. 2007; Masuyama and Watano 2010). Three of four samples identified as *Ceratopteris cornuta* showed relatively equal allelic proportions of 1:1 and 3:1, indicating a possible tetraploid; the fourth sample was ambiguous. Some samples of the known diploid *C. pteridoides* showed a similar pattern, with one putative diploid and three tetraploids. However, both species are known as a diploid so this may be a mis-assignment of ploidy by our analysis. *Ceratopteris gaudichaudii* had a high 1:1 allele ratio, suggesting that it is an allotetraploid. *Ceratopteris thalictroides* is also known to be a tetraploid (Lloyd 1974; Masuyama and Watano 2010), and individuals in our analysis were estimated to have 1:1, 2:1, and 3:1 allele ratios. Despite this variation, there was a distinct pattern between Old World and New World individuals: Old World *C. thalictroides* was most commonly allotetraploid (1:1 allele ratio), whereas New World *C. thalictroides* was either auto- or allotetraploid (relatively equal 1:1 and 3:1 allele ratios).

None of the individuals of *C. richardii* could be assigned to a distinct ploidy, due to the poor data quality (see Fig. 2). We had a few other samples with very poor quality data (see Supplementary data 1), and their ploidy assignments were almost all ambiguous. Mis-assignment of ploidy could be due to coverage being at the lower end of the scale needed for true ploidy assignments, but it also could indicate hybridization leading to genomic complexity across species. Including chromosome counts and flow cytometry data in future analyses would help better understand the potential variation in polyploidy origins of *C. thalictroides*.

## 4. Discussion

To the best of our knowledge, this is the first application of next-generation sequencing techniques to the complex genus *Ceratopteris*. Using -a population genomic, phylogenetic, and ploidy inferences, we recover similar species relationships found in previous studies, find support for *C. gaudichaudii* as genetically distinct, and identify a novel and distinct population from within *C. thalictroides sensu latu* in Central and South America.

### 4.1. Species boundaries in *Ceratopteris*

#### 4.1.1. *Ceratopteris cornuta* and *C. pteridoides*

*Ceratopteris cornuta* is found from Africa to Northern Australia, and *C. pteridoides* is endemic to the New World Tropics (Lloyd 1974). These two species are the most morphologically and genetically distinct in the genus. The sterile fronds of *Ceratopteris cornuta* are consistently deltoid to lanceolate in shape, and pinnate to bipinnate in dissection (Lloyd 1974). Our population genomic analyses distinguished all specimens of *C. cornuta* as a distinct population (at *K* greater than 4, (Fig. 3). All samples of *C. cornuta* from Africa group together at *K* = 2 - 7, along with one sample of *C. thalictroides* from Nepal *(Fraser-Jenkins 1564*) and an unnamed sample from Brazil *(L. Camargo de Abreu 21*). The sample from Nepal has a deltoid to lanceolate pinna shape, characteristic of *C. cornuta*. Lloyd (1974) notes that *C. cornuta* is known to hybridize with *C. thalictroides* in the eastern part of its range (Africa to India), however, the descriptions of such hybrids do not match the morphology this specimen (ie. *Blanford 500* NY from Sikkim). The unnamed sample from Brazil grouped with *C. cornuta* in both our population genomic and phylogenetic analyses. It has a very unusual morphology: the sterile leaves are 2-3 pinnate with very long-lanceolate pinnae. This could be a potential hybrid, or an undescribed morphotype of *C. thalictroides* in South America.

*Ceratopteris pteridoides* has perhaps the most unique morphology of the genus: sterile fronds are simple, palmately or pinnately lobed, with enlarged, air-filled stipes (petioles). They are most commonly found unrooted and floating on the surface of slow-moving water (Lloyd 1974). *Ceratopteris pteridoides* is also very strongly differentiated by the STRUCTURE analysis, with samples identified as *C. pteridoides* showing little to no introgression from other populations. Influence from *C. pteridoides* does appear in *C. cornuta*, *C. gaudichaudii*, and some samples of *C. thalictroides*. However, this may be an artifact of slightly lower sequencing coverage in these individuals.

#### 4.1.2. Cryptic species of *C. thalictroides*

There was one named cryptic species of *C. thalictroides* included in our study, *Ceratopteris gaudichaudii*, which has two varieties: *C. g. gaudichaudii* and *C. g. vulgaris* (Masuyama and Watano 2010); both were included in our analysis. *Ceratopteris g. vulgaris* has a much broader distribution than Guam endemic *C. g. gaudichaudii*: it can be found throughout the Pacific in Japan, the Philippines, Hawaii, Guam, Taiwan, Southeast Asia, and Northern Australia (Masuyama and Watano 2010). Individuals named as *C. thalictroides* from Taiwan, Hawaii, Japan, China, and Australia were identified via our population genomic and phylogenetic analyses as being *C. gaudichaudii*, although it is difficult to tell from our limited sampling which sub-species they align with. The individual from Australia was examined and keyed out morphologically to *C. g. vulgaris*, using the key from Masuyama and Watano (2010). This same individual was growing a few meters away from an individual of Old World *C. thalictroides*, indicating that these two morphologically similar species are growing sympatrically but with limited gene flow.

All of the samples from Hawaii included in this study show alignment to *C. gaudichaudii*. Wagner (1950) hypothesized that *Ceratopteris* was not native to Hawaii, and may have been introduced from Asia either as a food source or accidentally as a weed in taro patches. Lloyd (1973) noted that Hawaiian *Ceratopteris* was very similar to Japanese plants; the presence of *C. gaudichaudii* in Hawaii fits with this theory because the type specimen is from Japan. In addition, work by Hickok (1979) supported Hawaiian *Ceratopteris* as a distinct species from *C. thalictroides*. One sample from Hawaii was shown to be a potential hybrid between *C. gaudichaudii* and *C. thalictroides*. The specimen *(L. M. Crago 2005-058* US) is highly dissected and has may proliferous buds, which could indicate a hybrid or perhaps just an older plant.

Two named cryptic species of *C. thalictroides* were not included in our analysis: *C. oblongiloba* and *C. froseii. Ceratopteris oblongiloba* is found throughout Cambodia, Indonesia (Sumatra and Java), the Philippines (Luzon), and Thailand (Malay) (Masuyama and Watano 2010). Our only specimen from this range is from the Philippines, and aligns entirely with Old World *C. thalictroides*. This either indicates that we did not unknowingly sample *C. oblongiloba*, or it has a lesser degree of genomic distinction than the other cryptic species, *C. gaudichaudii*. *Ceratopteris froesii* is a Brazilian endemic characterized by very small fronds: about 4 cm or less for fertile leaves (Masuyama and Watano 2010). We only had four accessions from Brazil, and none were small plants. Inclusion of this unique species in future studies may help us further understand New World diversity of *Ceratopteris*.

#### 4.1.3. Ceratopteris thalictroides, sensu latu

Individuals of *C. thalictroides* that did not group with *C. gaudichaudii* break in to two groups: an Old World and a New World clade. These two clades are separated at all values of *K* in our STRUCTURE analysis (Fig. 3) and are paraphyletic (Fig. 4), indicating strong genetic, if not morphological, separation.

The Old World *C. thalictroides* clade includes mostly individuals from Asia and Australia, but also a few from South America. Increasing *K* in STRUCTURE did not break up this large group, suggesting that there is recent gene flow across the Pacific via long distance dispersal or human-mediated transport. One individual from Hawaii shows shared ancestry with *C. gaudichaudii* and Old World *C. thalictroides*, indicating that trans-Pacific gene flow might be possible. Another individual from Japan shows a similar pattern, but with a much larger proportion of its genome comprised of the Old World *C. thalictroides* population. There may be continued gene flow not only within Old World *C. thalictroides*, but between *C. thalictroides* and *C. gaudichaudii*.

New World *C. thalictroides* is strongly differentiated from Old World *C. thalictroides* in our population structure analysis, and shows virtually no gene flow between any other species at *K* greater than 3. This strong distinction between Old and New World is present in the phylogeny as well: New World *C. thalictroides* is placed in a clade as sister to *C. gaudichaudii*, and this clade is in turn sister to Old World *C. thalictroides*.

### 4.2. Independent polyploid origins of cryptic species

There is evidence from the present study and previous molecular work to support independent origins of cryptic species in *Ceratopteris thalictroides*. Adjie et al. (2007) identified Asian *C. thalictroides sensu latu* as paraphyletic and hypothesized that *C. thalictroides sensu strictu*, and the cryptic species *C. gaudichaudii* and *C. oblongiloba* are allopolyploid hybrids between *C. cornuta* and an unnamed, potentially extinct, diploid progenitor. Interestingly, we do not see evidence of extensive, recent admixture within species of *Ceratopteris* in our population structure analysis. There is some potential, limited hybridization within *C. cornuta* and a few accessions of *C. thalictroides*; but, *C. gaudichaudii*, Old World and New World *C. thalictroides* appear to be distinct species (Fig. 3). We did not include the Asian cryptic species *C. oblongiloba*, and so cannot say whether or not it is of recent hybrid origin.

Whereas our population structure analysis did not recover any evidence of recent hybridization in the *Ceratopteris thalictroides* species complex, we cannot rule out the possibility of more ancient hybridization. Our ploidy estimation indicates potential differences in polyploid origin between cryptic species of *C. thalictroides*: the Old World clade may be of allopolyploid origin, while the New World clade is of autopolyploid origin (Fig. 2, 4). *Ceratopteris gaudichaudii* is also potentially of allopolyploid origin, which supports the theory proposed by Adjie *et al*. (2007). Different polyploid origins of *C. thalictroides* were also suggested by McGrath *et al*. (1994) from their work on gene copy number across the genus. New World and Old World *C. thalictroides* are quite strongly differentiated by population structure and phylogenetic analyses (Fig. 3, 4), therefore separate polyploid origins for these two species could explain the mechanism of speciation. All of this evidence points to New World *C. thalictroides* being a distinct cryptic species of possible autopolyploid origin.

### 4.3. Evolutionary drivers in *Ceratopteris*

One of the most confusing aspects of *Ceratopteris* is the disparity between morphological and genetic distinction between species. The three diploid species are relatively morphologically distinct (Lloyd 1974), especially in comparison to the cryptic species of *C. thalictroides*. However, all the diploid species in the genus are known to hybridize with one another (Hickok and Klekowski 1974; Hickok 1977; Lloyd 1974). In contrast, *C. thalictroides* and associated cryptic species are much more similar morphologically, yet different populations (some named as cryptic species) can be almost entirely reproductively isolated (Hickok 1979; Masuyama and Watano 2010). These inconsistencies have contributed to the taxonomic confusion in the genus. Our phylogenetic reconstruction does, however, support multiple distinct species in the genus, despite known hybridization.

Weak reproductive boundaries are a theme across ferns, evidenced by numerous reticulate species complexes *(e.g.*, Barrington et al. 1989; Paris et al. 1989; Sessa et al. 2012), and intergeneric hybridization across 60 MY of evolution (Rothfels et al. 2015). Although ferns do have some pre- and post-zygotic barriers (Haufler et al. 2016), they are not as strong as in angiosperms (eg. Nettancourt 1997; Lafon-Placette and Köhler 2016). In addition, fern spores are very easily dispersed (Tryon 1970; Smith 1972), which limits isolation by distance. All of these factors may be limiting genetic diversification and enabling hybridization in *Ceratopteris*, despite its pan-tropical distribution. Continued gene flow may be contributing to the morphological similarities in the group, but there are certainly other factors at play.

Another potential explanation for morphological stasis in *Ceratopteris* relies on the influence ecological pressures (Bickford et al. 2007). For a fern, *Ceratopteris* has an unusually short life cycle of only about four months from spore to spore-bearing adult (Stein 1971). It grows in temporary water sources such as pond edges, swamps, or tarot patches (Lloyd 1974). Because of its ephemeral habitat, there are likely selective pressures for maintaining morphological stasis and rapid generation times. Fossil spore and leaf evidence also support morphological stasis in the Ceratopteridoideae dating back about 47 MY (Dettmann and Clifford 1992; Rozefelds et al. 2016). Fossil leaf impressions of the extinct genus *Tecaropteris* from Australia are very similar morphologically to extant *Ceratopteris* (Rozefelds et al. 2016). Australia has become increasingly dry and seasonal since the Eocene, when *Tecaropteris* lived (Rozefelds et al. 2016; McKenna et al. 2010). Today, *Ceratopteris* is exceedingly difficult to find during the dry season in northeastern Australia, but plentiful during the wet season, even growing as a weed in garden ponds (Dr. Ashley Field, personal communication). Almost all leaves of *Ceratopteris* species are very thin, delicate, and herbaceous (Lloyd 1974). When removed from the plant they wilt into an unrecognizable state within minutes (personal observation, SPK). These low-energy investment leaves can be grown quickly and without an excess of resources from the plant (Reich 2014; Wright et al. 2004), potentially as a result of the fast life cycle employed by *Ceratopteris*; any major shift in leaf development might inhibit *Ceratopteris* from proceeding through its life cycle before its water source dries up. Therefore, despite the genetic diversity in the genus, morphological diversity has changed very little due to niche constraints.

## 5. Conclusion

In summary, our findings suggest that there may be more cryptic species of *Ceratopteris thalictroides* yet to be discovered, especially in the New World. The potential for cryptic species in the Neotropics has been noted by several authors (Masuyama and Watano 2010; Lloyd 1974), and this area of the world remains one of the least studied parts of the range of *Ceratopteris*. Considering that this is where the model species *C. richardii* is native, more work on all species of the genus in the New World is especially warranted. Researching *Ceratopteris* in its natural environment is critical to inform future lab studies on *C. richardii*, and the potential incorporation of other species for comparative studies. Most importantly, understanding speciation and cryptic species within *Ceratopteris* is important for its use as model system, as well as increasing our knowledge of evolutionary processes in ferns and across vascular land plants.

## Acknowledgements

The authors thank the University of Wisconsin Biotechnology Center DNA Sequencing Facility for providing DNA extraction, library prep, and DNA sequencing facilities and services. We would also like to thank the University of Utah Center for High-Performance Computing, particularly Anita Orendt, for providing computational resources for data analyses. Thank you to Ashley Field and Tzu-Tong Kao helping with field work; Jacob S. Suissa and David Barrington for sending herbarium specimens; and Ryan Choi and Jacob S. Suissa for their thoughtful comments and edits to the manuscript. SPK is funded by a National Science Foundation Graduate Research Fellowship and the Joseph E. Greaves Endowed Scholarship from Utah State University. WDP and the Pearse lab are funded by National Science Foundation ABI1759965, NSF EF1802605 and United States Department of Agriculture Forest Service agreement 18CS11046000041.

## Appendix A. Supplementary material

Supplementary data 1.

